# Figeno: multi-region genomic figures with long-read support

**DOI:** 10.1101/2024.04.22.590500

**Authors:** Etienne Sollier, Jessica Heilmann, Clarissa Gerhäuser, Michael Scherer, Christoph Plass, Pavlo Lutsik

## Abstract

**Summary:** The vast amount of publicly available genomic data requires analysis and visualization tools. Here, we present figeno, an application for generating publication-quality FIgures for GENOmics. Figeno particularly focuses on multi-region views across genomic breakpoints and on long reads with base modifications. Additionally, we support epigenomic data including ATAC-seq, ChIP-seq or HiC, as well as whole genome sequencing data with copy numbers and structural variants.

**Availability and Implementation:** Figeno is available as a python package with both a command line and graphical user interface. It can be installed via PyPI and the source code is available at https://github.com/CompEpigen/figeno.

**Supplementary information:** Supplementary data is provided.

## 1 Introduction

Recent advances in sequencing have led to the generation of many different types of genomic and epigenomic data, which require visualization tools for facilitating analysis and interpretation. An important requirement for such tools is the support of various data types, for large datasets, and for generating publication-quality images. The integrative genomics viewer (IGV)^1^ enables interactive exploration of genomics data, but lacks support for some data types like chromatin conformation (e.g. HiC). Furthermore, IGV has limited export capabilities, making it subpar for generating publication-quality figures. Other tools are geared towards a specific data type, e.g., trackplot^2^ for bigwig files, while others like PyGenomeTracks^3^ can handle more data types. An important limitation of most genomics visualization software is that they only display one genomic region at a time, which is admittedly sufficient for many use cases but for example precludes the visualization of novel chromatin interactions across breakpoints in HiC data. NeoLoopFinder^4^ has a visualization component which enables the visualization of HiC data across breakpoints, but it is not primarily a visualization tool. Third generation sequencing technologies (Oxford Nanopore Technologies, Pacific Biosciences) provide long reads which can be phased to each parental haplotype, as well as base modification information. Most visualization software do not support long reads, while new tools such as methplotlib^5^ and methylartist^6^ have been specifically designed to display base modifications from long reads, but lack other common features such as support for bigwig and HiC data. Here, we introduce figeno, an application for visualizing genomics and epigenomics data, including long reads, which allows multiple regions to be visualized simultaneously.

## 2 Figeno: a flexible, versatile and user-friendly solution for genomic plotting

### 2.1 Figeno overview

Figeno is a visualization tool for genomic data tracks (Figure 1A). Figeno’s key features are (i) the support of many data types; (ii) multi-region views; (iii) the support for base modifications from long reads; (iv) export into various image formats (see Supplementary Table 1 for a comparison to other visualization software). Of the ten types of tracks that can be visualized with figeno (Figure 1B), the first three tracks serve general purposes: “chr_axis” specifies the genomic coordinates, “genes” shows annotated genes in the region, and “bed” allows for additional custom genomic annotations. Gene files are provided for the human reference genomes hg19 and hg38, but users can also provide custom annotation files for further reference genomes (in NCBI RefSeq or GTF format). “bigwig” tracks allow for the visualization of epigenetic data types including ChIP-seq or ATAC-seq, and “hic” tracks can be used to visualize HiC data (.cool format). For alignment files in bam format, figeno visualizes either the alignments themselves, the coverage, or base modification frequency (see section long-read support). Finally, figeno supports the visualization of whole genome sequencing data with tracks for copy number and structural variants. The figures can be exported as bitmap (png) or vector (svg, pdf) graphics formats for direct use in publication or further processing with a vector graphics editor. A configuration file in json format serves as input to figeno. In addition to custom generation of the configuration file, we provide a graphical user interface (GUI) for interactive figure configuration (Supp. Fig. 1).

**Figure 1.**
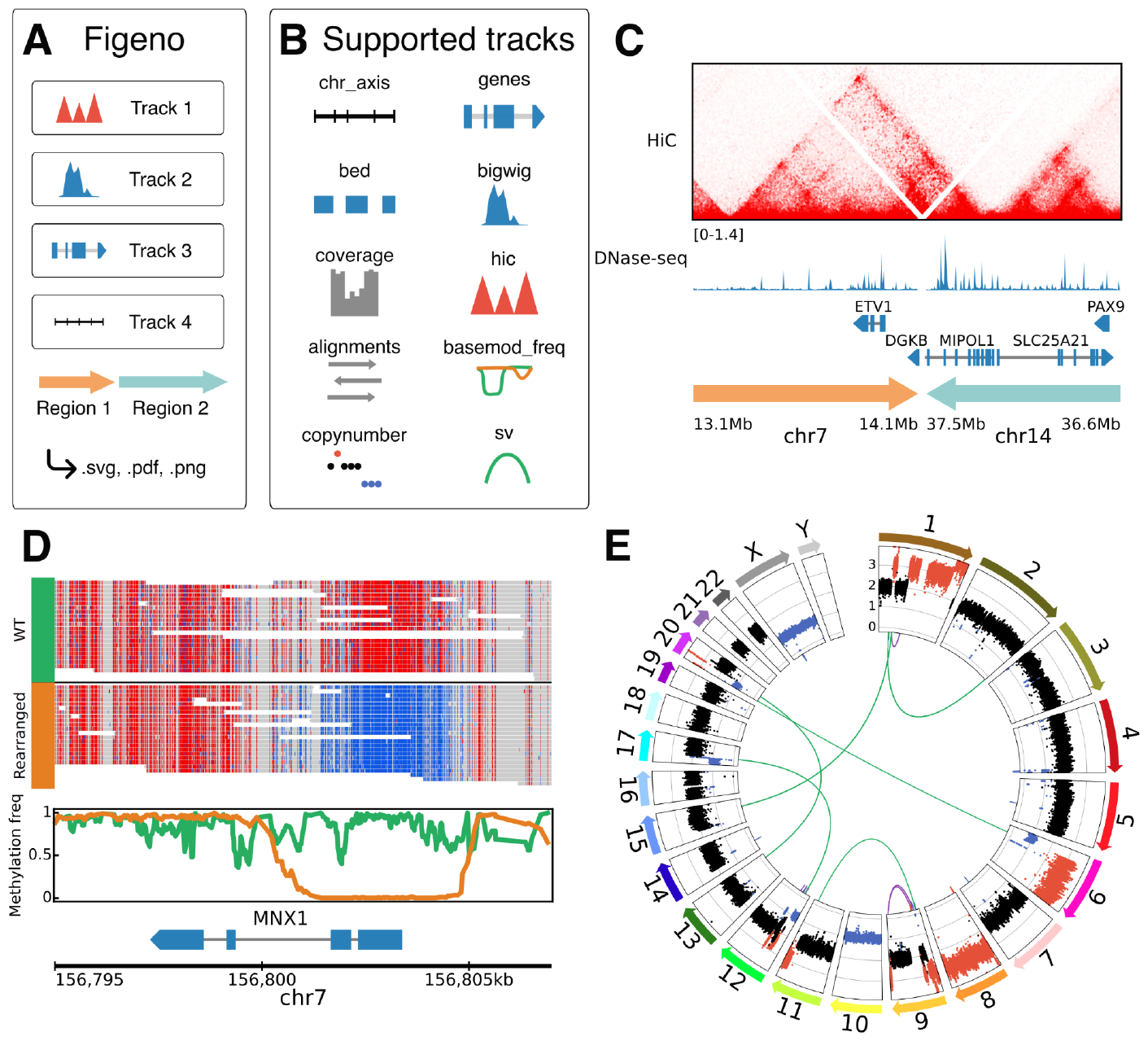
Figeno overview and example outputs. **A**. Overview of figeno. **B**. List of all 10 track types available within figeno. **C**. HiC data around a breakpoint in the LNCaP cell line^4^, and DNase-seq bigwig track from the LNCaP cell line, downloaded from the ENCODE project (ENCSR000EKT). **D**. Allele-specific DNA methylation at the *MNX1* locus in the GDM-1 cell line, based on nanopore sequencing data. For the alignments track, red indicates a methylated CpG site and blue unmethylated. **E**. Circular plot showing copy numbers and structural variants for the THP-1 cell line (data from the cancer cell line encyclopedia^10^). Copy-number gains are indicated in red and losses in blue.

### 2.2 Multi-region plotting

Figeno can plot one or multiple regions simultaneously. An important feature is that, although some tracks will be plotted independently for each region, other tracks can show interactions across regions: HiC tracks show chromatin interactions across genomic regions (Fig. 1C), structural variant (SV) tracks can show breakpoints across regions (Fig. 1E), alignment tracks can link multiple alignments from split reads aligned to different regions (Supp. Fig. 2), and scales in bigwig and coverage tracks can automatically be adjusted across regions.

### 2.3 Support for long-read data

Figeno implements features unique to long-read sequencing data, including grouping alignments by haplotype (if the HP tag is set in the bam file, for example by WhatsHap^11^) and coloring by base modification (if the MM and ML tags are provided in the bam file). In addition to the most common base modification (5mC), any base modification (e.g., 5hmC, 6mA) can be visualized. Furthermore, two base modifications can be visualized at the same time (e.g. 5mC and 5hmC). We also support a “basemod_freq” track to specify the base modification frequency at each position in a haplotype-aware manner. The alignments and “basemod_freq” tracks can display allele-specific DNA methylation. The leukemic cell line GDM-1 harbours a translocation between chromosomes six and seven, which activates *MNX1* by enhancer hijacking^12^. By visualizing allele-specific methylation in this region using figeno, we observed allele-specific methylation, where the wild type allele is methylated at the *MNX1* promoter while the rearranged allele is hypomethylated (Fig.1D). Finally, reads aligning to different genomic regions (“split-reads”) can be visualized by a line connecting all alignments from the same read (Supp. Fig. 2).

### 2.4 Customizable figure layouts

The default layout is “horizontal” and results in the regions being arranged horizontally from left to right. This layout is best-suited for most applications, but we also offer several other layouts that can be particularly useful for plotting whole genome sequencing data. First, the “circular” layout can be used to display all regions on a circle, for example to show all chromosomes in a circos plot (Fig.1E). For settings where we only want to look at 2-8 chromosomes with breakpoints between them, we also provide a novel “symmetrical” layout, where the regions are aligned on two rows, but the order of the tracks in the top row is reversed, which is particularly useful when an SV track is displayed between the two rows (Supp. Fig. 3).

## 3 Implementation and availability

Figeno is implemented in python and uses matplotlib^7^ for plotting. It relies on pysam^8^ for reading bam files, pybigwig for reading bigwig files, cooler^9^ for reading HiC data, and vcfpy for reading vcf files. The GUI was created using the javascript framework React, and utilizes a local Flask-based webserver. The time required to generate a figure depends on the number and types of tracks, on the size of the genomic regions being visualized and on the computer being used, but figeno generally only takes a couple of seconds to generate a figure. The code is completely open-source and is available on Github (http://github.com/CompEpigen/figeno) under the GPL-3 license.

## 4 Conclusion

Taken together, figeno is a rich and user-friendly visualization tool for genomics and epigenomics data, especially for structural rearrangements and long-read data. It supports an extensive collection of input formats (Supp. Table 1) and provides a GUI for enhanced usability by a broad range of users, from experienced bioinformaticians to beginners. To support them, we also provide extensive and detailed documentation on ReadTheDocs (https://figeno.readthedocs.io/).

## Acknowledgements

We thank Anna Riedel, Marvin Mayer, Simge Kelekçi, Yunhee Jeong and Katherine Kelly (Division Cancer Epigenomcs, DKFZ, Heidelberg, Germany) for testing figeno.We thank Alexander Vogel (Oxford Nanopore Technologies) for valuable discussion on setting up nanopore sequencing.

## Author contributions

ES developed figeno and wrote the original manuscript draft. JH performed nanopore sequencing of the GDM-1 cell line. CG, CP, MS and PL provided supervision. All authors reviewed and approved the final manuscript.

## Data generation and availability

ONT data was generated specifically for the manuscript. DNA from the GDM-1 cell line was extracted with the QIAamp DNA micro kit. Short fragments were removed with the PacBio SRE XS kit. Library preparation was performed with the ligation sequencing kit SQK-LSK114. The library was sequenced on one PromethION flow cell for 96h, with a wash and reload after 48h, which resulted in a coverage of 56x with an N50 of 21.5kb. Nanopore sequencing data for the AML cell line GDM-1 will be made available on the SRA (SRA ID: SRR28257102; the accession is kept private before publishing, reviewer link: https://dataview.ncbi.nlm.nih.gov/object/PRJNA1085236?reviewer=lb2hkvjnmo4rfoe6euh028srks).

All other data used in this manuscript were already publicly available: the HiC data of the LNCaP cell line from GEO accession GSE161493^4^, the DNase-seq data of LNCaP from the ENCODE project (file ID ENCSR000EKT), the WGS data of the THP-1 from the cancer cell line encyclopedia^10^ at the SRA with accession SRR8670675.

All data required to reproduce the figures in the manuscript are provided at https://github.com/CompEpigen/figeno/tree/main/test_data.

## References

1. Robinson, J. T. et al. Integrative genomics viewer. Nat. Biotechnol. 29, 24–26 (2011).

2. Mayakonda, A. & Westermann, F. Trackplot: a fast and lightweight R script for epigenomic enrichment plots. Bioinforma. Adv. vbae031 (2024) doi:10.1093/bioadv/vbae031.

3. Lopez-Delisle, L. et al. pyGenomeTracks: reproducible plots for multivariate genomic datasets. Bioinformatics 37, 422–423 (2021).

4. Wang, X. et al. Genome-wide detection of enhancer-hijacking events from chromatin interaction data in rearranged genomes. Nat. Methods 18, 661–668 (2021).

5. De Coster, W., Stovner, E. B. & Strazisar, M. Methplotlib: analysis of modified nucleotides from nanopore sequencing. Bioinformatics 36, 3236–3238 (2020).

6. Cheetham, S. W., Kindlova, M. & Ewing, A. D. Methylartist: tools for visualizing modified bases from nanopore sequence data. Bioinformatics 38, 3109–3112 (2022).

7. Hunter, J. D. Matplotlib: A 2D Graphics Environment. Comput. Sci. Eng. 9, 90–95 (2007).

8. Li, H. et al. The Sequence Alignment/Map format and SAMtools. Bioinformatics 25, 2078–2079 (2009).

9. Abdennur, N. & Mirny, L. A. Cooler: scalable storage for Hi-C data and other genomically labeled arrays. Bioinformatics 36, 311–316 (2020).

10. Ghandi, M. et al. Next-generation characterization of the Cancer Cell Line Encyclopedia. Nature 569, 503–508 (2019).

11. Martin, M. et al. WhatsHap: fast and accurate read-based phasing. 085050 Preprint at 10.1101/085050 (2016).

12. Weichenhan, D. et al. Translocation t(6;7) in AML-M4 cell line GDM-1 results in MNX1 activation through enhancer-hijacking. Leukemia 1–4 (2023) doi:10.1038/s41375-023-01865-5.

