## Supplementary material for "Figeno: multi-region genomic figures with long-read support"

**Supplementary Table 1**: Feature comparison of genomic visualization software in comparison to figeno.

| | Figeno | IGV | Trackplot | PyGenome Tracks | NeoLoop Finder | Methplotlib | Methylartist |
|---|---|---|---|---|---|---|---|
| Programming language | Python | Java | R | Python | Python | Python | Python |
| Multi-region support | ✔ | ✔ | ✘ | ✘ | ✔ | ✘ | ✘ |
| Bigwig | ✔ | ✔ | ✔ | ✔ | ✔ | ✘ | ✘ |
| HiC | ✔ | ✘ | ✘ | ✔ | ✔ | ✘ | ✘ |
| Reads with base modification | ✔ | ✔ | ✘ | ✘ | ✘ | ✔ | ✔ |
| Base modification frequency | ✔ | ✘ | ✘ | ✘ | ✘ | ✔ | ✔ |
| WGS (copy number, SV) | ✔ | ✘ | ✘ | ✘ | ✘ | ✘ | ✘ |
| Graphical user interface | ✔ | ✔ | ✘ | ✘ | ✘ | ✘ | ✘ |
| Interactive exploration | ✘ | ✔ | ✘ | ✘ | ✘ | ✘ | ✘ |
| Vector graphics export | ✔ | —* | ✔ | ✔ | ✔ | ✔ | ✔ |

*IGV can export to svg, but the resulting files are usually very large and difficult to edit further.

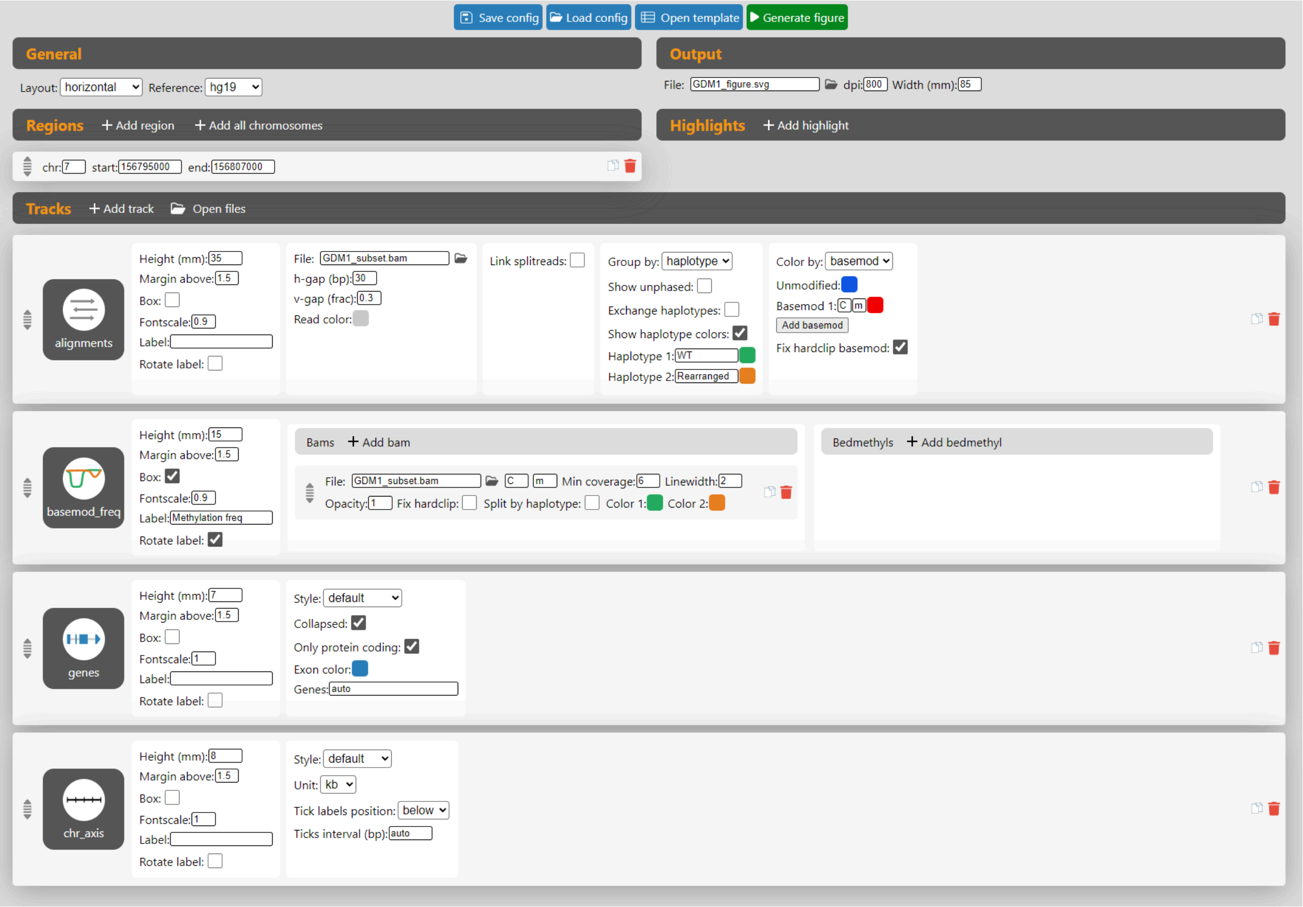


**Supplementary Figure 1: graphical user interface.** Screenshot of figeno's graphical user interface, here with the configuration for generating the allele-specific methylation plot for GDM-1 (Fig. 1D), with one region and four tracks: alignments, basemod_freq, genes and chr_axis.

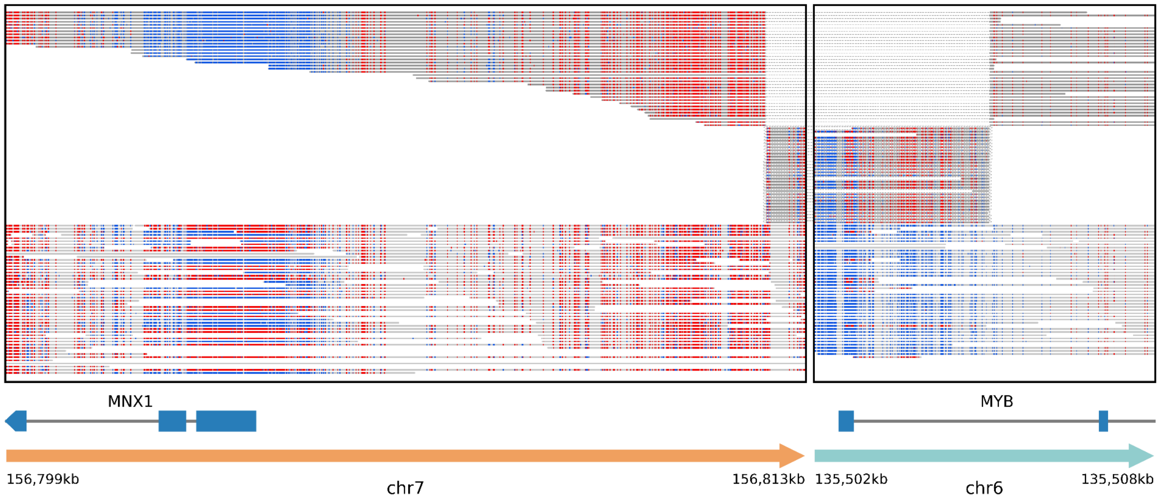


**Supplementary Figure 2: split-read visualization.** Nanopore reads for the GDM-1 cell line around the t(6;7) breakpoints. Alignments coming from the same read are linked together by dashed lines.

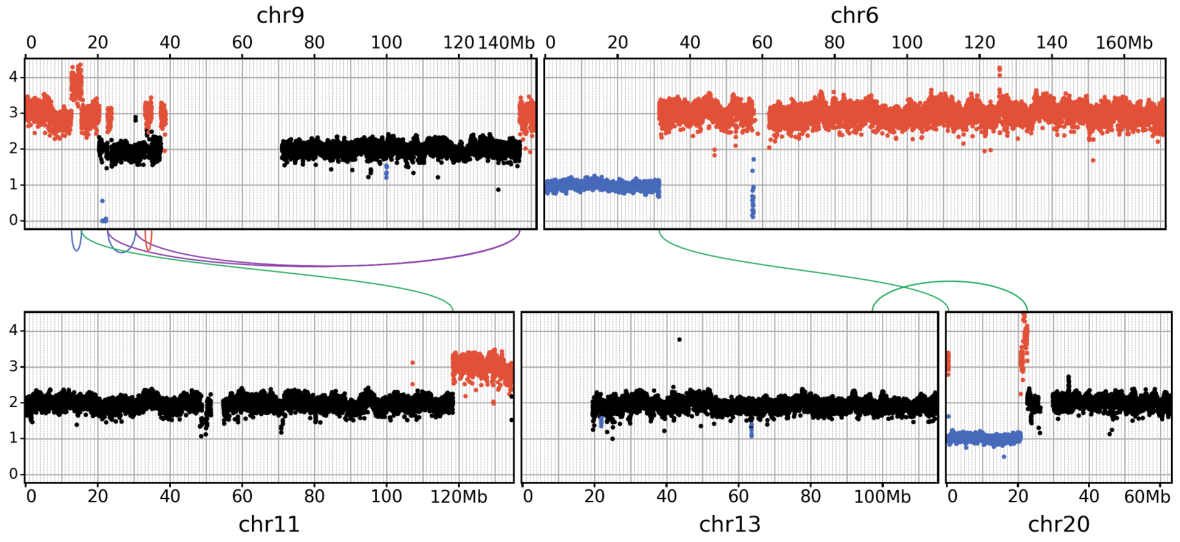


**Supplementary Figure 3: symmetrical layout.** Copy number and structural variants for chromosomes 6, 9, 11, 13, and 20 for the THP-1 cell line (whole genome sequencing data from the cancer cell line encyclopedia).
